# Impaired activation of Transposable Elements in SARS-CoV-2 infection

**DOI:** 10.1101/2021.02.25.432821

**Authors:** Matan Sorek, Eran Meshorer, Sharon Schlesinger

## Abstract

Transposable element (TE) transcription is induced in response to viral infections. TE induction triggers a robust and durable interferon (IFN) response, providing a host defense mechanism. Still, the connection between SARS-CoV-2 IFN response and TEs remains largely unknown. Here, we analyzed TE expression changes in response to SARS-CoV-2 infection in different human cellular models. We find that compared to other viruses, which cause global upregulation of TEs, SARS-CoV-2 infection results in a significantly milder TE response in both primary lung epithelial cells and in iPSC-derived lung alveolar type 2 cells. TE activation precedes, and correlates with, the induction of IFN-related genes, suggesting that the limited activation of TEs following SARS-CoV-2 infection may be the reason for the weak IFN response. Diminished TE activation was not observed in lung cancer cell lines with very high viral load. Moreover, we identify two variables which explain most of the observed diverseness in immune responses: basal expression levels of TEs in the pre-infected cells, and the viral load. Finally, analyzing the SARS-CoV-2 interactome, as well as the epigenetic landscape around the TEs that are activated following infection, we identify SARS-CoV-2 interacting proteins, which may regulate chromatin structure and TE transcription in response to a high viral load. This work provides a functional explanation for SARS-CoV-2’s success in its fight against the host immune system, and suggests that TEs could be used as sensors and serve as potential drug targets for COVID-19.

**Key points:**

- Unlike other viruses, SARS-CoV-2 invokes a weak and inefficient transposable element (TE) response
- TE induction precedes and predicts IFN response
- Basal TE expression and viral load explain immune responses
- Distinct chromatin and enhancer binding factors occupancy on TEs induced by SARS-CoV-2

## Introduction

Coronaviruses are a diverse group of single-stranded positive-strand RNA viruses infecting a wide range of vertebrate hosts. These viruses are thought to generally cause mild upper respiratory tract illnesses in humans such as the common cold. However, infection with severe acute respiratory syndrome-related coronavirus 2 (SARS-CoV-2), which causes Coronavirus Disease-2019 (COVID-19), can result in a cytokine storm, which develops into acute respiratory distress syndrome and acute lung injury, often leading to reduction of lung function and even death^1^. The fatality of SARS-CoV-2 increases substantially with age^2^. Unlike other airborne viruses, SARS-CoV-2 is unusually effective at evading the early innate immune responses, such as type I and type III interferons (IFN-I; IFN-III). This is partially achieved by viral proteins that antagonize various steps of dsRNA-activated early host responses^3,4^. However, the understanding of the kinetics of IFN response in mild and severe COVID-19 patients is lacking.

First, Blanco-Melo et al. assessed the transcriptional response to SARS-CoV-2 infection in different cellular models^1^ and found that SARS-CoV-2 does not elicit robust IFN expression in lung epithelial cells, while multiple pro-inflammatory cytokines were highly expressed. Second, Huang et al. assessed the transcriptional response to SARS-CoV-2 infection in alveolar type 2 cells (iAT2s) derived from induced pluripotent stem cells (iPSCs) by *in vitro* differentiation^5^. The authors found that SARS-CoV-2 infection in these cells results in an inflammatory phenotype with an activation of the NF-κB pathway, and a delayed IFN signaling response. Other recently published data support these conclusions^6,7^, although the outcome of IFN response on the virus and its host are not clear^8^.

Transposable elements (TEs) are abundant sequences in the mammalian genome that contain multiple regulatory elements and can amplify in a short evolutionary timescale^9^. Lately, it was found that TEs induction stimulate antiviral response, via both *cis* and *trans* mechanisms^10^. First, TEs have coopted to shape the transcriptional network underlying the IFN response, and some TEs serve as enhancers of antiviral genes in diverse mammalian genomes^11,12^. TEs are also enriched in enhancers of CD8+ T lymphocytes-specific genes, suggesting that their upregulation might influence not only the innate but also the adaptive, immune response^13^. What’s more, due to the similarities between TEs and viral transcripts, cells sometimes misidentify them as invading viruses and trigger the innate immune nucleic acid sensors (e.g. RIG-I, MDA-5 and cGAS) controlling IFN response. Consequently, genome-wide global induction of TEs is common during some viral infections in humans, and transposon induction is part of the first wave response to viral infection^14,15^. For example, infection by highly pathogenic avian influenza viruses elicits TEs induction^16^. As a result, TE dsRNA is formed, recognized by the host sensors and activates both the NFkB and the IFN pathways, thus enhancing immune response^17^. Collectively, these findings suggest a causative link between TE induction and the intensity of the IFN response^18^.

This link is also evident in aging as TE basal expression increases with age^19,20^. Consequently, aging is associated with sterile inflammation which include erroneous IFN response, or the ‘Inflammaging’ phenomena^21^. On the other hand, at the molecular level, aging is associated with decreased heterochromatin-associated marks, e.g. H3K9me3^22^ and DNA methylation^23^. Interestingly, early antiviral IFN responses are impaired and delayed in aged individuals, resulting in increased risk of COVID-19 complication^24^. Similarly, elevated TE expression is found in autoimmune pathologies like arthritis and systemic lupus erythematosus^18,25^, which are suggested as risk factors for severe COVID-19^26^.

Although some studies have addressed TEs expression following SARS-CoV-2 infection^27,28^, none has considered the cell type and basal TE expression levels as important characteristics of their analysis. Here, we suggest that SARS-CoV-2 infected cells that fail to activate an immediate and effective TE response will be more likely to demonstrate a late immune response. Notably, in primary cells, the IFN response to SARS-CoV-2 is observed only 96 hours after infection, unlike that observed in cell lines^29^. This correlation between the IFN response and TE expression levels strengthens our model and gives rise to a hypothesis that link between mild TE levels and the ineffective innate response to SARS-CoV-2 infection in some individuals^29^. Given the high correlation between TEs activation and higher IFN response, we suggest a possible use for TEs in COVID-19 prognosis

## Results

### TE induction in response to SARS-CoV-2 infection is limited in normal lung cells

Since TEs are induced in response to many viral infections^14^, we first analyzed recently published datasets of primary human lung epithelial cells (NHBE) infected with influenza (IAV)^30^ or SARS-CoV-2^1^. NHBE cells mimic infected human lung cells, showing cytopathic effects after SARS-CoV-2 infection^31^. In these cells IAV infection caused, as expected, a global increase in TE subfamilies expression across all TE families, but SARS-CoV-2 did not (Figure 1A). As expected, IFNβ treated cells also activated TE expression levels (Figure 1A) because many TEs have IFN-responsive sequences and are upregulated following the induction of IFN response^25,32^. This raised an intriguing hypothesis that SARS-CoV-2 may avoid the robust TE expression response that often follows a viral infection.

**Figure 1.**
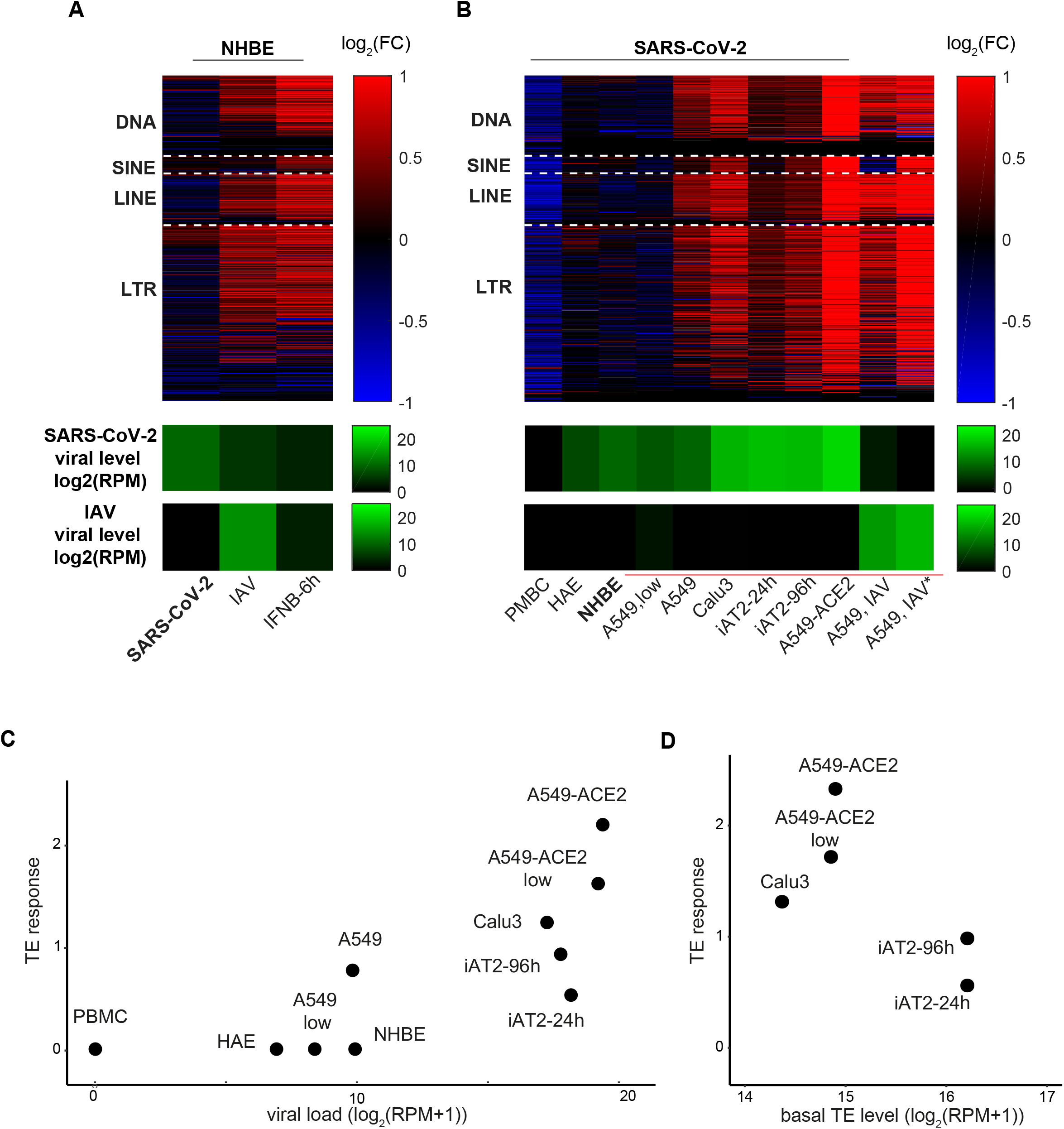
TEs expression changes in response to SARS-CoV-2 and IAV infections. **A.** Log2-fold change in expression level of TE subfamilies (DNA, SINE, LINE and LTR) in NHBE cells in response to IFNB treatment and in response to SARS-CoV-2 and IAV infections. SARS-CoV-2 viral levels (green) are depicted in the bottom panel. **B.** Same as (A) for SARS-CoV-2 infection in different cellular systems and IAV infection in A549 cell line. **C-D.** TE induction levels are correlated with SARS-CoV-2 viral levels (C) and also with TE basal levels pre-infection (D). Linear regression coefficients are 0.17 and -0.39 for viral load and basal TE level, respectively (R^2^=0.78). PBMC were removed from the regression because they had essentially zero viral load.

To examine this hypothesis, we expanded our data analysis to more cell types infected with SARS-CoV-2 and other viruses. For SARS-CoV-2, we added lung cancer cell lines A549 and Calu3, and iAT2 primary cells, which represent a young state of normal lung cells, as well as primary human airway epithelial (HAE) cells and peripheral blood mononuclear cells (PBMCs)^5^. As expected, we found that other viruses induced a marked activation of TE (Figure S1A). Importantly, we observed that the viral load affects the strength of the TE response to SARS-CoV-2 infection: the higher the viral load the stronger the TE upregulation (Figure 1B, note green heat map at the bottom denoting the viral load of each sample).

Remarkably, although viral load explains much of the observed TE response (Figure 1C, R^2=0.66, p=0.0078. See Methods for details), incorporating the TE basal levels into the model improved the accuracy of the prediction of the TE response (Figure 1D, R^2=0.78, p-value=0.0095). This is because for both low viral load and high viral load, the TE induction levels in the primary cells, which have a higher TE basal level, were mild compared to the transformed cell lines (Figure 1B). A549 cells had a stronger TE response compared to the primary NHBE cells although both had similar low viral load levels, while at higher viral load levels Calu3 and A549 cells expressing ACE2 – the virus entry receptor - had a stronger TE response compared to the iAT2 cells. This pattern was consistent among the different TE classes (Figure S1A). Therefore, viral load is not solely responsible for the upregulation of TEs expression. The other two contributing factors are the identity of the virus, where SARS-CoV-2 induces weaker activation than IAV and other viruses (Figure S1A), and the identity of the cells, where in primary cells that have a higher basal TE level the TEs are less induced compared with transformed cell lines.

### Upregulation of TEs is positively correlated with high IFN response

Because IFN expression had previously been associated with TE expression^11,17,33^, we next tested the relationship between IFN and TEs expression during SARS-CoV-2 infection. In agreement with the more robust TE induction, we found a significantly larger number of upregulated IFN genes (Table S1) in the cancer cell lines one day after infection with SARS-CoV-2 compared with primary cells, including NHBE cells (Figure 2A). Importantly, NHBE cells did respond to IAV infection and to IFNβ treatment by a significant induction of IFN genes (Figure 2B and Figure S1B) and TE expression levels (Figure 1A) already after 4-12 hours, excluding the possibility that these cells simply fail to activate TEs or induce IFN response. While infection of iAT2 cells with SARS-CoV-2 resulted in a mild but detectable TE response (Figure 1B, larger TE log fold change [LFC] compared to NHBE cells, p< 1e-6 Kolmogorov-Smirnov test) one day post infection, it showed almost no IFN response, even weaker than infected NHBE cells (smaller LFC of IFN genes, p<0.0001 Kolmogorov-Smirnov test, Figure 2A). This mild TE response slightly increased after 4 days and was accompanied by a significant IFN response (Figure 2A). This suggests that the early TE response precedes the late IFN response in these cells.

**Figure 2.**
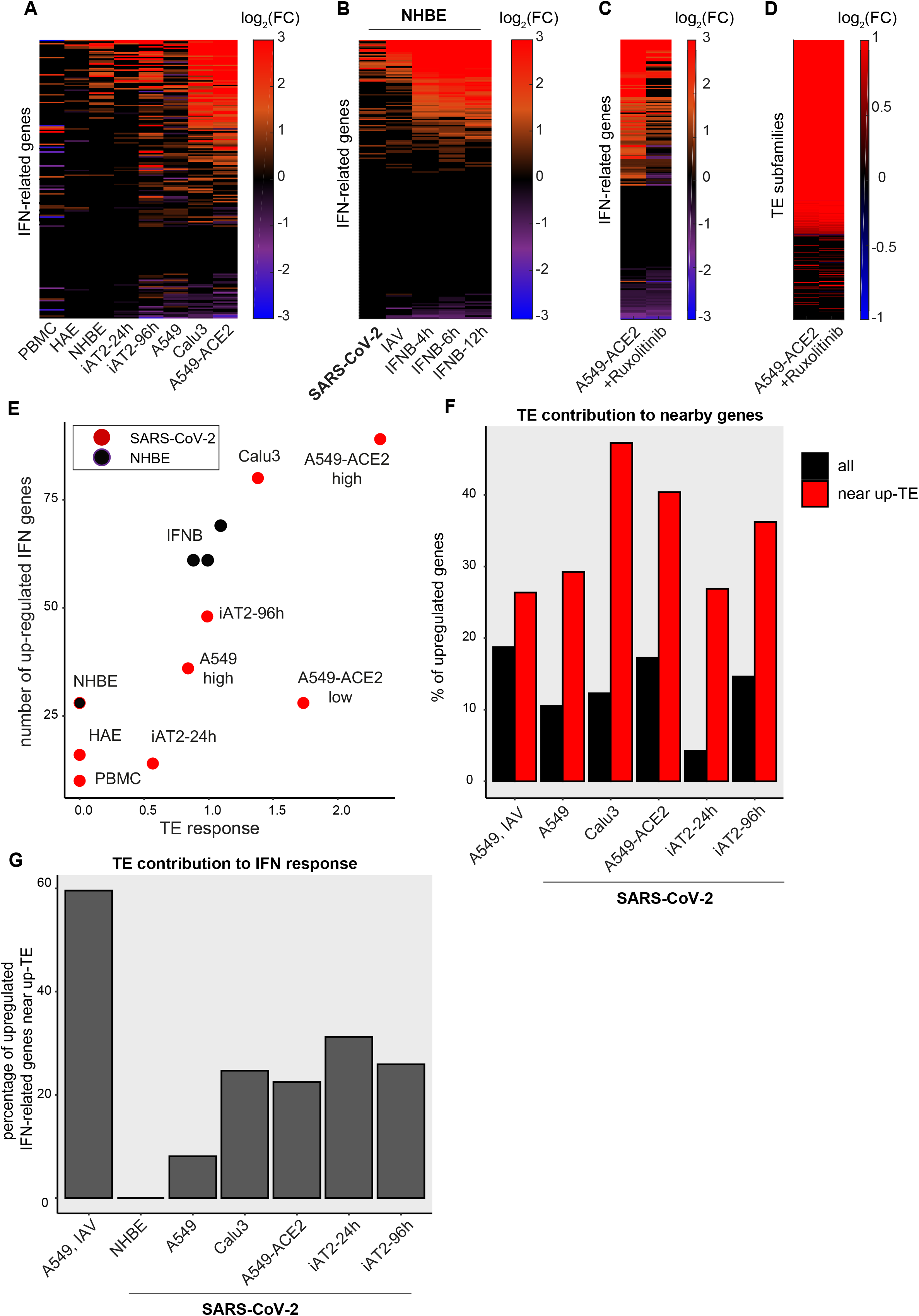
TE induction precedes and predicts IFN response. **A.** The IFN transcriptional response for SARS-CoV-2 infection in difference cellular systems. **B.** Same as (A) for NHBE and iAT2 cells. For A549 IAV there is one experiment from Blanco-Melo et al. and one from Schmidt et al. (marked by an asterisk). **C.** The IFN transcriptional response of A549 cells over-expressing the ACE2 receptor dramatically decreases upon Ruxolitinib treatment. **D.** TE up-regulation persists even with Ruxolitinib treatment. **E.** IFN transcriptional changes correlate with the TE induction levels among SARS-CoV-2-infected cells (red) and among NHBE cells (purple). TE response is calculated as the 95-percentile of log2 fold-change of all TE subfamilies. **F.** The percentages of upregulated genes out of all genes and the upregulated genes that are near upregulated TEs in response to SARS-CoV-2 infection in different cellular systems. **G.** The fraction of upregulated IFN-related genes that are located near upregulated TEs.

The idea that TE activation is inducing IFN response following SARS-CoV-2 infection was also supported by A549 cells over-expressing the ACE2 receptor that were infected with SARS-CoV-2. These cells, when treated with Ruxolitinib, a JAK1/2 inhibitor that is known to reduce inflammatory response^34^, showed reduced IFN response (Figure 2C), while the global TE response remain high (Figure 2D). This shows that although IFN response upregulate TE expression, TE overexpression is not solely dependent on the IFN response. Finally, although done on different tissues, genetic backgrounds and multiplicity of infection (M.O.Is), the magnitude of IFN changes strongly correlated with TE expression changes (Figure 2E). This correlation (p<0.05, permutation test on Spearman correlation) was specific to IFN related genes (and epifactors, see below), as other random groups of genes were not correlated with TE changes. These, and other^14,32^ results, indicate that while IFNβ treatment activate TEs expression, TEs upregulation is an early response to viral infection, which precede the IFN response and induce it in a positive feedback loop^14^.

Since TEs were shown to function as regulatory elements, or enhancers, for adjacent host genes encoding critical innate immune factors^11^, we also tested the relation between expression changes in individual TEs and their neighboring genes. In general, genes that were adjacent to upregulated TEs, were prone to be upregulated as well (Figure 2F). Focusing on IFN-related genes (Table S2) located near up-regulated TEs revealed an even stronger effect, suggesting that this IFN gene induction by an adjacent TE occurs also following SARS-CoV-2 infection (Figure S1C). However, while ~60% of the upregulated IFN-related genes were located near upregulated TEs in response to IAV infection, only 10-30% of the upregulated IFN related genes were located near upregulated TEs in response to SARS-CoV-2 infection (Figure 2G). This suggests that although following SARS-CoV-2 infection TEs have the capacity to serve as *cis*-regulatory enhancers to nearby genes including IFN related genes, the TEs nearby IFN related genes remain mostly silent. In concordance, gene ontology (GO) analysis revealed that while the upregulated genes adjacent to upregulated TEs in response to IAV infection were enriched for cytokine receptor binding genes, no immune related function was enriched among those in response to SARS-CoV-2 infection.

Taken together, these data suggest that while TE induction following IAV infection precedes and contributes to the expression of IFN-related genes^30^, SARS-CoV-2 infection fails to upregulate the TEs coopted for immune activation. According to this hypothesis, TE induction is a crucial step in the activation of the anti-viral immune response against RNA viruses^35^. However, TE response to SARS-CoV-2 infection is limited: considerably less TEs are activated, the level of their upregulation is lower, and most importantly, their specificity is altered: those TEs that induce immune response are not the ones activated by SARS-CoV-2 infection.

### The epigenetic signature of SARS-CoV-2 induced TEs

Regulation of TEs transcription is largely achieved through epigenetic silencing^36^. To understand the nature of the distinctive TE regulation of those TEs that do go up following SARS-CoV-2 infection, we investigated which specific histone modifications are found on the upregulated TEs before SARS-CoV-2 infection (Table S2).

To this end, we used a large-scale dataset including multiple ChIP-seq profiles for histone modifications (HMs) in uninfected A549 cells^37^. Since for A549 cells we also have pre- and post-SARS-CoV-2 infection data, it allowed us to examine the ‘epigenetic signature’^38^ (relative enrichment of HMs) around the TEs that are upregulated post-infection of SARS-CoV-2 and IAV (Figure 3–5, Figure S2–S3 and Table S3).

**Figure 3.**
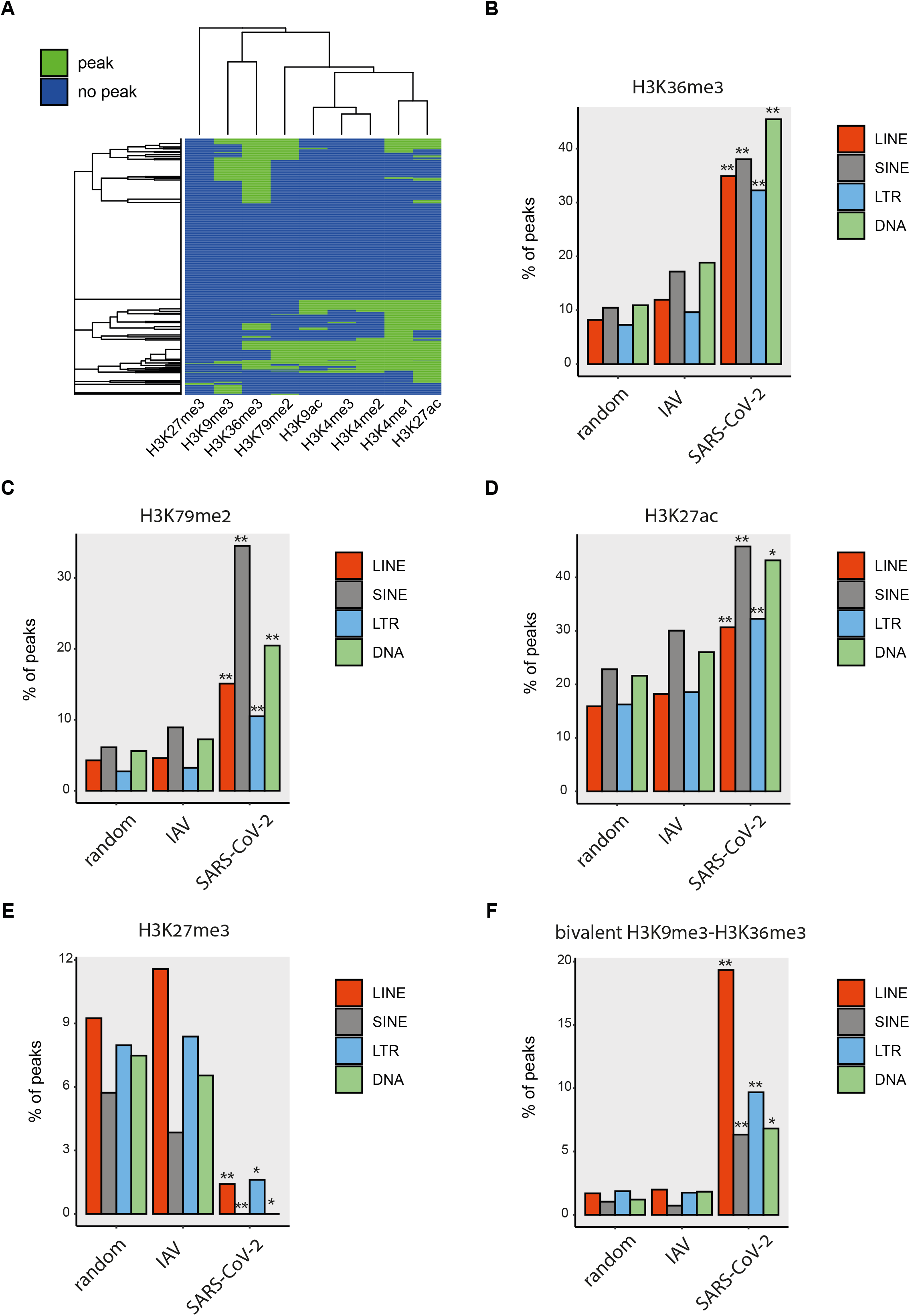
TE classes that are induced by SARS-CoV-2 in A549 cells have a unique epigenetic profile. **A.** Hierarchical clustering of histone modifications signal in non-infected A549 cells around all upregulated TEs in response to SARS-CoV-2 infection in A549 cells. **B-E.** Percentage of TEs with peaks of H3K36me3 (B), H3K79me2 (C), H3K27ac (D), H3K27me3 (E) on SARS-CoV-2 induced TEs, IAV induced TEs and on all expressed TEs outside genes. **F.** Same as B-E for a bivalent signature of H3K9me3 and H3K36me3 mark. Asterisks mark significance level of the difference compared to all expressed TE outside genes: one asterisk marks FDR-adjusted p-value < 0.05 and two asterisks mark p < 0.001, requiring at least a 2-fold difference in Fisher’s exact test.

We found that the TEs that were upregulated in response to SARS-CoV-2 infection in A549 cells of all different classes were enriched for active histone marks in uninfected cells, with a subset of TEs marked by H3K36me3 as well as the combination of H3K27ac, H3K4me3, H3K79me2 and H3K9ac (Figure 3A–E and Figure S2). This was consistent among different ChIP-seq experiments for the same histone modification (Figure S3). SINEs and DNA elements upregulated in response to SARS-CoV-2 infection were especially enriched for active marks compared to both random TEs and IAV-induced TEs. By contrast, LINEs that were upregulated in response to SARS-CoV-2 infection were highly enriched for a bivalent signature of the repressive H3K9me3 mark together with the active H3K36me3 mark (Figure 3G and Figure 4). Interestingly, all classes of SARS-CoV-2-induced TEs were depleted for H3K27me3 in the uninfected cells (Figure 3F). This is in stark contrast to IAV-induced TEs, which show no such depletion (Figure 3F).

**Figure 4.**
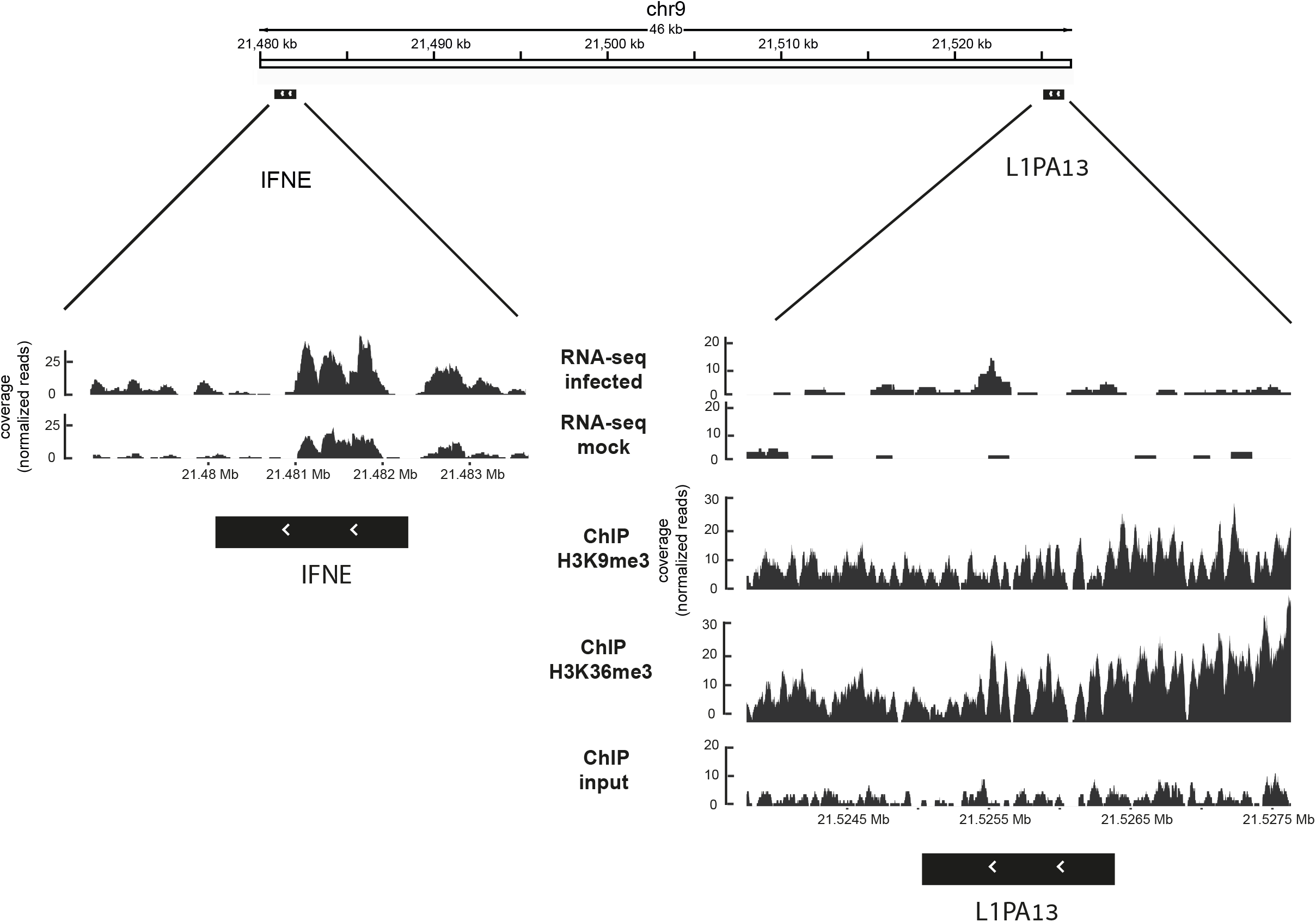
up-regulated TE with unique epigenetic signature contributes to gene expression response to SARS-CoV-2 infection. Shown are UCSC tracks for the H3K9me3-H3K36me3 bivalently marked LINE copy L1PA13, which is up-regulated following SARS-CoV-2 infection in A549 cells. The TE is located near the IFN-related gene IFNE which is also up-regulated in these cells in response to SARS-CoV-2 infection.

Repeating the analysis at the TE family level, we found that the largest enrichment was in the specific L1 and L2 LINE families and in the MIR and Alu SINE families (Figure 5). These families showed a strong enrichment especially in H3K36me3 and H3K79me2, and were depleted for H3K27me3. The H3K36me3 and H3K79me2 were also significantly enriched on the main ERV/LTR families. Overall, these results suggest that the TEs that are upregulated in response to SARS-CoV-2 in A549 cells have a distinct epigenetic profile, which differs from that of TEs upregulated by IAV, as well as from the general epigenetic profile of TEs in the human genome. Specifically, SARS-CoV-2-induced TEs are devoid of H3K27me3, and are comprised of two major subsets of TEs: *i*. SINEs and DNA elements marked by a highly active chromatin profile, and *ii*. a bivalent group of LINEs marked by both repressive and active marks, which keeps them in a poised state ready for infection-induced expression. This attests at the relative failure of the SARS-CoV-2 infected cells to activate TEs in a repressive chromatin state as IAV does.

**Figure 5.**
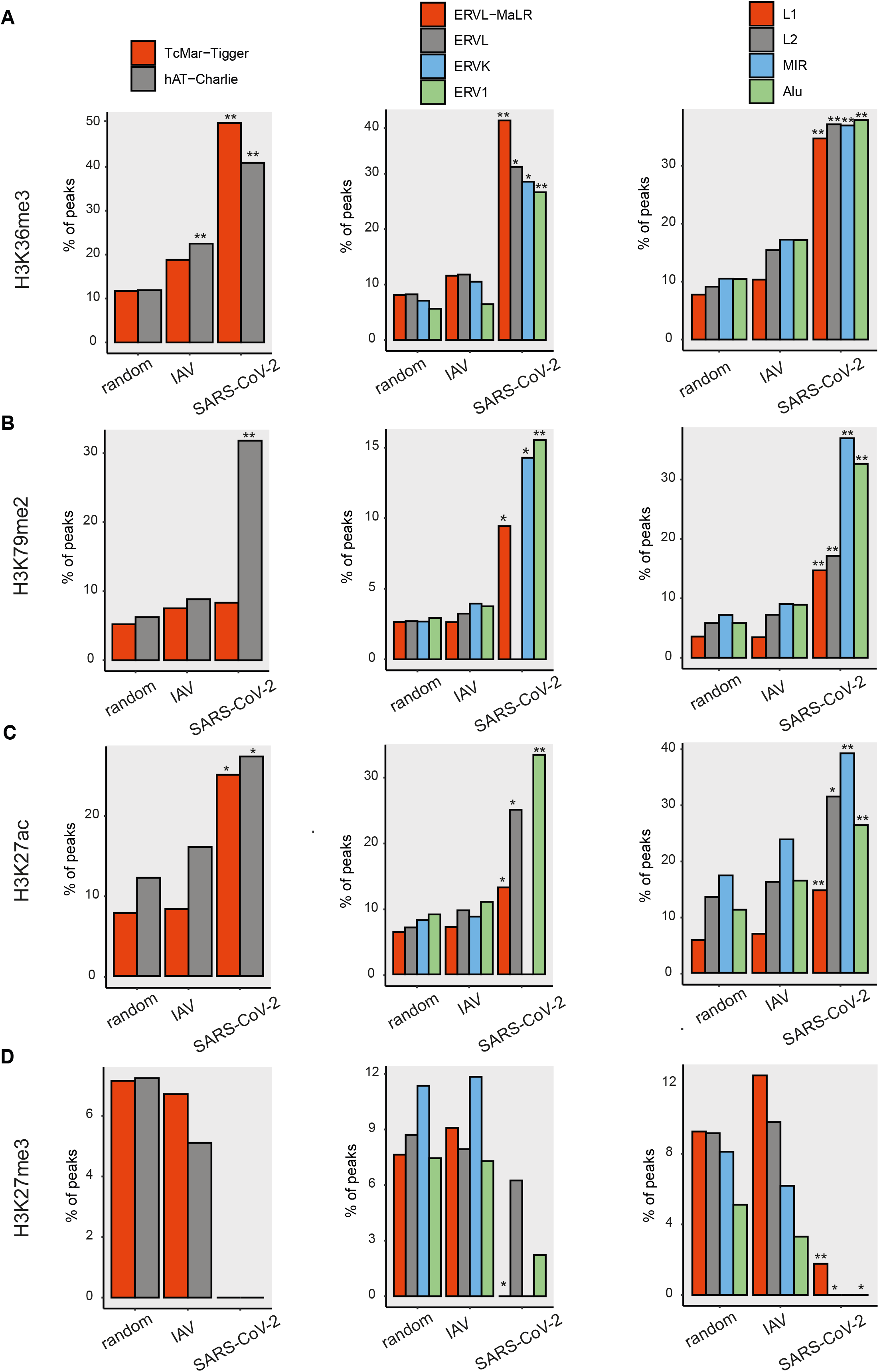
TE families that are induced by SARS-CoV-2 in A549 cells have a unique epigenetic profile. **A.** Percentage of TEs with peaks of H3K36me3, **B.** H3K79me2, **C.** H3K27ac, **D.** H3K27me3 on SARS-CoV-2 induced TEs, IAV induced TEs and on all expressed TEs outside genes. Shown are DNA families (first column), LTR families (second column) and LINE and SINE families (third column). Asterisks mark significance level of the difference compared to all expressed TE outside genes: one asterisk marks FDR-adjusted p-value < 0.05 and two asterisks mark p < 0.001, requiring at least a 2-fold difference in Fisher’s exact test.

### The transcriptional signature of infected cells

Seeking a possible mechanism for the activation of TEs with this distinctive chromatin modification pattern, we searched for genes, expression of which changes in correlation with TE response. We focused on genes that had a high correlation with TE response both in the SARS-CoV-2 infection from Blanco-Melo et al. and in the iAT2 infected cells from Huang et al. We found that the inversely correlated genes were enriched for mitochondrial-related genes and processes (Figure 6A and Table S4, and see Methods), consistent with previous reports^39^. By contrast, genes that were positively correlated with TE response among all samples were enriched, in addition to type I interferon production, for chromatin, DNA and enhancer binding, RNA Pol-II binding, transcription factor and cofactor binding as well as histone binding, demonstrating a clear epigenetic and chromatin-related signature (Figure 6A and Table S4).

**Figure 6.**
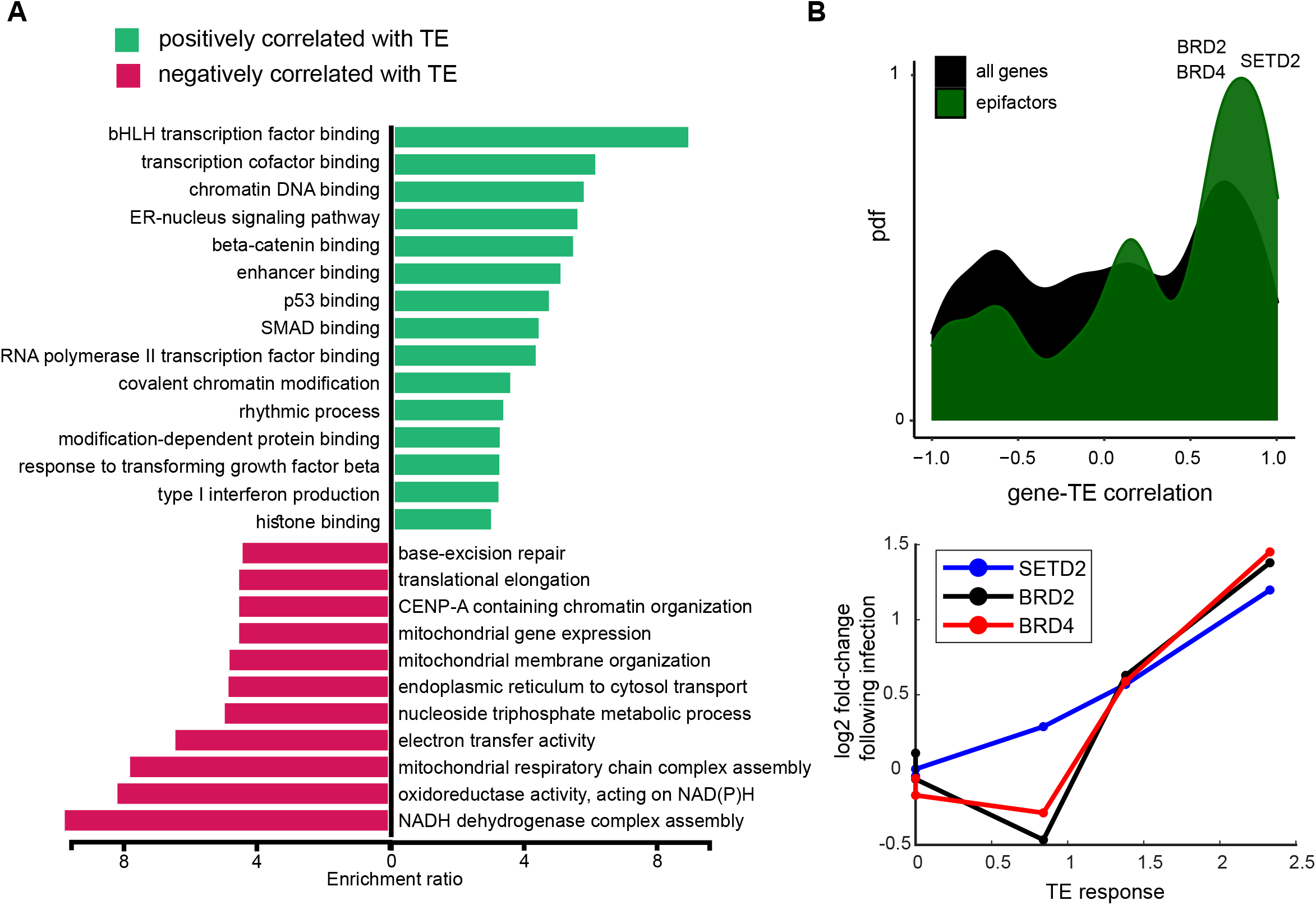
TE response is correlated with epigenetic and mitochondrial gene expression changes. **A.** Gene ontology analysis of genes correlated (green) and inversely-correlated (red) with TE response to SARS-CoV-2 infection. **B.** Distribution of TE response-gene correlations for genes that increase with TE expression in iAT2 cells. Black represents correlation distribution of all genes. Correlation distribution of epifactors are in green. **C.** The LFC of gene expression following SARS-CoV-2 infection vs the TE response in different systems for SETD2, BRD2 and BERD4, three SARS-CoV-2 interactome-related epigenetic factors.

Indeed, intersecting the positively correlated genes with all genes that encode for chromatin binding proteins, or epifactors^40^, showed highly significant enrichment (Figure 6B, green. 44 genes, fold-enrichment=2.33, p < 10^−7^, Fisher’s exact test). Interestingly, SETD2, the human H3K36 lysine trimethylase, was among the epifactors that were positively correlated with TE response (Figure 6C). H3K36me3 marks gene bodies of active genes. In addition, SETD2 methylation of STAT1 is crucial for interferon response^41^ and its H3K36 methylation contributes to ISG activation, pointing at its role in the cellular response to viral infection. Finally, recent evidence shows that SETD2 is essential for microsatellite stability, implicating its role in non-genic transcriptional regulation^42,43^. Interestingly, SETD2 is also among the interacting proteins of SARS-CoV-2, and compared with other interactomes of coronaviruses as well as of different IAV strains, we found that SETD2 is specific for SARS-CoV-2 (Table S5).

We therefore searched for more epifactors in the SARS-CoV-2 interacting proteins. Among the ten epifactors that interact with SARS-CoV-2, we found the histone acetylation related proteins BRD2, BRD4, which were highly correlated ti the TE response (Figure 6C), and HDAC2, all of which are, once again, SARS-CoV-2-specific (Table S5). We also found two TE-related epifactors: the SARS-CoV-2-specific interacting epifactor DDX21, a DNA damage and dsRNA sensing protein, and MOV10, an RNA helicase that also restricts LINE expression^44^, which interacts with SARS-CoV-2, as well as with IAV. These observations suggest that SARS-CoV-2 may affect TE expression through interaction with a subset of specific epifactors.

## Discussion

In this study, we re-analyzed published data to examine the link between SARS-CoV-2 infection and transcriptional activation of Transposable Elements (TEs). We find that in normal lung epithelial cells, SARS-CoV-2 does not induce global upregulation of TEs, as observed for IAV and other RNA viruses (Figure 7). This phenomenon is in correlation with the viral load and with the intensity of IFN response in the infected cells. The low IFN response after infection is also correlated with high basal TE expression in the uninfected cells^21,45^. Interestingly, a recent large-scale study of TE expression in different tissues during aging, demonstrated that TE expression levels are gradually elevated in most tissues as a function of age^20^.

**Figure 7.**
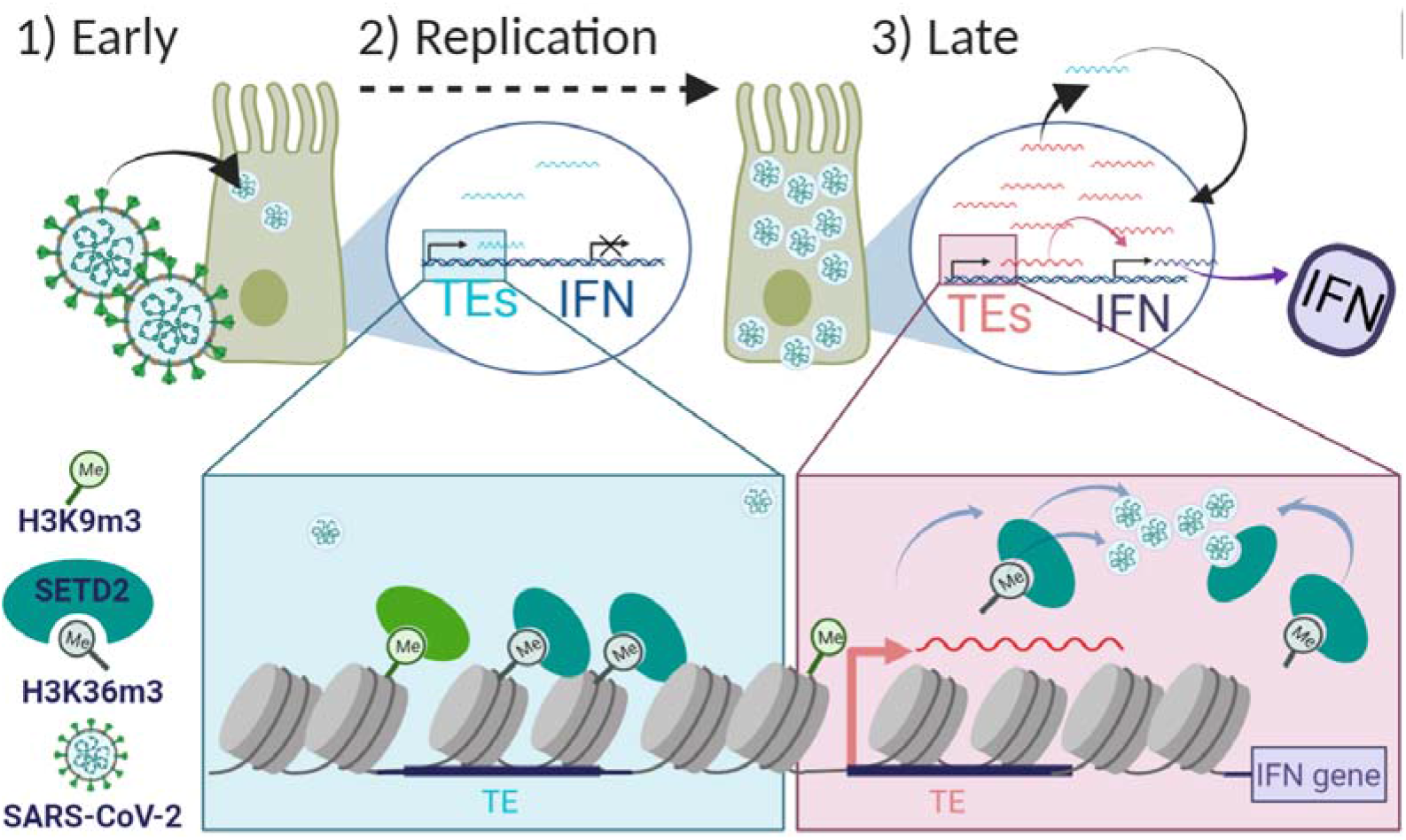
A model for SARS-CoV-2 impaired TE activation. Early following SARS-CoV-2 infections of normal lung epithelial cells, viral load is low and IFN response is still absent. SETD2 is normally localized on the H3K36me3 enriched chromatin. Next, viral replication (2) increases viral load and induces TE activation and IFN response (3). The hypothesis (in the pink box) is that specific SARS-CoV-2 interacting proteins sequester chromatin factors, such as SETD2, away from the H3K36me3 enriched TEs, which are then transcribed, inducing IFN response that might be too weak or too late for the host defense

Since TE expression rises with age^19,20^, and age is the most significant risk factor for COVID19-related death, we hypothesize that high basal level of TE expression desensitizes the TE induction response to viral infection, explaining the age-related decline in survival. In contrast, young people should benefit from higher SARS-CoV-2 induced TE overexpression that, in turn, prompts IFN response early in the disease course^6^. Our “TE desensitization” model makes several predictions. First, it anticipates that induction of TEs would precede the IFN response. Second, the cellular TE activation response to SARS-CoV-2 should be associated with basal levels of TE expression, where lower basal levels predict stronger cellular responses. As such, and thirdly, it anticipates that in older cells, which would have a higher basal level of TEs and would be “TE desensitized”, the TE activation response will be significantly milder. This would lead, fourthly, to a less effective IFN induction response, allowing SARS-CoV-2 to operate “under the radar” at early disease stages when the viral load is still low. Finally, at later stages as the viral load accumulates, selected TEs are induced. Those TEs are characterized by histone modifications, regulators of which are found to interact with the SARS-CoV-2 proteome.

As our model predicts, we showed that SARS-CoV-2 elicits a weaker TE activation response compared with other viruses, and a weaker TE activation response in primary cells compared with cancer cells. These primary cells have higher initial levels of TEs. We speculate that in old normal cells, low TE expression fails to induce viral mimicry and thus fails to initiate early innate immune reaction. This is in line with previously published data that link TE expression to immune reaction^17,33^, as well as studies that show suppression of dsRNA-activated early host responses following SARS-CoV-2 infection^3,4^. Our own analysis also shows a highly significant correlation between TE activation and the induction of IFN response.

Cells that showed a large TE response also had a strong IFN response to SARS-CoV-2 infection, but not through adjacent upregulated TEs. This is because the immune response can be induced by other mechanisms beside by TEs acting as enhancers for immune genes^36^. For example, the TE themselves can also act in *trans*, being sensed as dsDNA and dsRNA by the cells sensors for these species Therefore, the impaired TE activation following SARS-CoV-2 infection may have a double impact, yielding the delayed IFN response seen in COVID-19 patients.

In addition, SARS-CoV-2 viral load seems to be linked with TE expression level changes. Accordingly, we were able to compare TEs induced following a rigorous SARS-CoV-2 infection to that of IAV infection in A549 cells. Interestingly, the TEs that are upregulated in response to SARS-CoV-2 in A549 cells showed a distinct epigenetic profile, which differed from that of TEs upregulated by IAV, as well as from the general epigenetic profile of TEs in the human genome. Specifically, SARS-CoV-2-induced TEs are devoid of H3K27me3 and enriched for H3K36me3, a subset of which is bivalently marked also with H3K9me3. This atypical pattern was identified as a mark for poised enhancers that control surrounding gene expression^46^. Here, the activation is potentially mediated by the H3K36 trimethylase SETD2. SETD2 expression is closely correlated with TE induction, and it specifically interacts with the SARS-CoV-2 NSP9 protein.

Together, these data suggest a model where SARS-CoV-2 entry modifies SETD2 deposition or activity, leading to aberrant H3K36me3 enrichment on a subset of TEs. Consequently, these TEs are transcribed, leading to the induction of the IFN response. In older age, where basal TE levels are high, the changes in chromatin structure and histone modifications will have minor effects, either because the TEs are promiscuously expressed or because of the compromised TE-induced IFN response.

Overall, by re-analyzing data published by Huang et al., Blanco-Melo et al. and others, we provide evidence that unlike other viruses, which strongly induce TEs following infection^14^, SARS-CoV-2 infection has a relatively weak effect on TE expression in primary lung epithelia. One potential consequence of this reduced TE induction in normal cells is weak activation of the IFN response which usually responds to TEs via a positive feedback loop mechanism^47^. Our analysis further showed that SARS-CoV-2 could elicit a strong TE induction in lung cancer cells, Calu3 and A549. This could potentially be explained by epigenetic aberrations often observed in transformed cells, rendering them prone to TEs dysregulation^17,33,47^.

## Methods

### Datasets

The original sequencing datasets for IAV infection in A549 cells and the Blanco-Melo et al datasets can be found on the NCBI Gene Expression Omnibus (GEO) server under the accession numbers GSE133329 and GSE147507, respectively. The data produced by Huang et al. can be found at GSE153277. The PBMC and HAE data can be found at the Genome Sequence Archive in BIG Data Center under the accession number CRA002390 and on GEO under the accession number GSE153970, respectively.

### Preprocessing and alignment

Raw reads from GSE147507 and GSE133329 were trimmed to remove Illumina adapters using the Trimmomatic software^48^ version 0.39. We used the STAR aligner version 2.7.1a^49^ to align raw reads to human RefSeq reference genome (GRCh38). We used the parameters --outFilterMultimapNmax 100 and --winAnchorMultimapNmax 200 to allow for large numbers of multi-mapped reads for downstream analysis of TEs. For ChIP-Seq data, alignment with STAR was followed by filtering out of non-uniquely mapped reads.

### TE and gene expression quantification

Gene expression was analyzed separately. Genes were considered significantly up-(down-) regulated if they had at least 1.5 fold difference and FDR corrected p-value < 0.05. We quantified TE expression changes both at the level of TE subfamilies and at the level of individual repeat loci. For TE subfamily quantification we used TEtranscripts from the TEToolKit^50^. The TEtranscript algorithm quantifies TE subfamilies and genes simultaneously by assigning together multi-mapped reads which are associated with the same TE subfamily. Human repeat annotations for hg38 were downloaded from TEtranscripts site. TEtranscripts was run using --mode multi and -n TC. This was followed by differential expression analysis using DESeq2^51^. TE response was quantified using the 95-percentile of the log2 fold-change of all TE subfamilies.

To quantify gene expression and to determine the locations of individual TEs that change in expression we used featureCounts v2.0.0^50,52^ from the Subread package which uses only uniquely mapped reads. Simple repeat elements were removed prior to the analysis. The TE and gene count matrices were combined, followed by DE-Seq2 to compare between mock and infected cells. Individual TEs were considered significantly up-(down-) regulated if they had at least 1.5 fold difference and p-value < 0.05. In order to robustly calculate the IFN response, we used all genes associated with the following GO terms: GO_0035458, GO_0035457, GO 0035456, GO_0035455, GO_0034340, as well as genes associated with the following pathways in pathcards (https://pathcards.genecards.org/Pathway): Immune response IFN alpha/beta signaling super-pathway and pathways 2747, 2388, 213 (Table S1). IFN-related genes were considered significantly up-(down-) regulated if they had at least 1.5 fold difference and p-value < 0.05. The list of epifactors was downloaded from https://epifactors.autosome.ru/^40^.

### ChIP-Seq signal quantification

To calculate the ChIP profile of different histone modification on TEs, we used the processed output files from the ENCODE project, which are filtered for the ENCODE blacklist regions (see Table 1). For each TE we defined the flanking region as the TE location and its surrounding 500 bp up- and down-stream. If the flanking region intersection with ChIP peak locations was non-empty then this TE was considered as TE with peak. For histone modification clustering (Figure 3 and Figure S2) we used the Jaccard metric. Enrichment of peaks on up-regulated TEs was calculated using hypergeometric test. Genome tracks were produced using the ggbio package in R^53^.

**Table 1:** ChIP-Seq data from ENCODE project^37^:

| ChIP Name<br>(in this<br>manuscript) | Experiment<br>identifier<br>(ENCODE) |
| --- | --- |
| H3K27Ac | ENCFF443MVY<br>ENCFF918FET<br>ENCFF747IZX |
| H3K9ac | ENCFF350BNS |
| H3K79me2 | ENCFF705EEZ |
| H3K36me3 | ENCFF584FSY |
| H3K4me3 | ENCFF832EOL<br>ENCFF361TGO<br>ENCFF535EYL |
| H3K9me3 | ENCFF287WGC |
|  | ENCFF164FDB |
| H3K27me3 | ENCFF684ZZH |
| H3K4me2 | ENCFF536TYP<br>ENCFF794FQH |
| H3K4me1 | ENCFF116HIQ<br>ENCFF594YDK |

### TE-gene correlation analysis

For TE response-gene correlation, we used Spearman rank correlation between the 95-percentile TE subfamily log fold-change and the log fold-change of genes in the SARS-CoV-2 infected cells. We used NHBE, A549, Calu3, and A549-ACE2 cells infected using M.O.I.=2 in addition to A549 cells infected with M.O.I.=0.2. Genes were defined as highly correlated and inversely-correlated if the correlated was larger than 0.97 and smaller than -0.97 among these samples, respectively, and if, in addition, the direction of change aligned with that in the iAT2 cell between d1 and d4 after infection. These cutoffs corresponded to approximately the top and bottom 5%.

### Interactome analysis

For interactome analysis, we downloaded publically available interactome lists (see Table 2). A gene was considered in SARS-CoV-2 interactome if it was included in at least 3 out of 5 sources, and as a SARS-CoV-1 interacting partner if it was included in both relevant sources.

**Table 2:** interactome sources

| virus | Interactome papers |
| --- | --- |
| SARS-CoV-2 | 54-58 |
| MERS-Cov | 57 |
| SARS-CoV-1 | 54,57 |
| HCoVNL63 | 54 |
| HCoV229E | 54 |
| IAV (different strains) | 59 |
| EBV | 60 |
| WNV | 61 |

## Figure legends

**Figure S1: impaired TE activation in response to SARS-CoV-2 infection. A.** The 95-percentile of TE subfamilies log2 fold changes in response to IFNB treatment, and to SARS-CoV-2 and IAV infection in different cellular systems by class. **B.** The distribution of log2 fold-changes of TE subfamilies in response to SARS-CoV-2 infection and IFNB treatment at different times in NHBE cells. **C.** The percentage of up-regulated IFN-related genes located near TEs that are up-regulated following SARS-CoV-2 and IAV infections out of all IFN-related genes located near up-regulated TEs.

**Figure S2: epigenetic signature of up-regulated TEs in response to SARS-CoV-2 and IAV infections compared to all expressed TEs. A-B.** Hierarchical clustering of histone modifications signal in non-infected A549 cells around all up-regulated TEs in response to IAV infection (A) and 5000 random expressed TEs (B) in A549 cells. **C-F.** Percentage of TEs with peaks of H3K9Ac (C), H3K4me3 (D), H3K4me2 (E), H3K4me1 (F) on SARS-CoV-2 induced TEs, IAV induced TEs and on all expressed TEs outside genes.

**Figure S3: epigenetic profile of SARS-CoV-2 up-regulated TEs is consistent among different experiments. A-F.** Percentage of TEs with peaks additional H3K27ac (A-B), H3K4me3 (C-D), H3K4me2 (E) and H3K4me1 (F) datasets on SARS-CoV-2 induced TEs, IAV induced TEs and on all expressed TEs outside genes.

## References

1 Blanco-Melo, D. et al. Imbalanced Host Response to SARS-CoV-2 Drives Development of COVID-19. Cell 181, 1036–1045 e1039, doi:10.1016/j.cell.2020.04.026 (2020).

2 Verity, R. et al. Estimates of the severity of coronavirus disease 2019: a model-based analysis. The Lancet infectious diseases 20, 669–677 (2020).

3 Yin, X. et al. MDA5 Governs the Innate Immune Response to SARS-CoV-2 in Lung Epithelial Cells. Cell Rep 34, 108628, doi:10.1016/j.celrep.2020.108628 (2021).

4 Ancar, R. et al. Physiologic RNA targets and refined sequence specificity of coronavirus EndoU. RNA 26, 1976–1999, doi:10.1261/rna.076604.120 (2020).

5 Huang, J. et al. SARS-CoV-2 Infection of Pluripotent Stem Cell-Derived Human Lung Alveolar Type 2 Cells Elicits a Rapid Epithelial-Intrinsic Inflammatory Response. Cell Stem Cell Sep 18, S1934–5909, doi:10.1016/j.stem.2020.09.013 (2020).

6 Park, A. & Iwasaki, A. Type I and Type III Interferons - Induction, Signaling, Evasion, and Application to Combat COVID-19. Cell Host Microbe 27, 870–878, doi:10.1016/j.chom.2020.05.008 (2020).

7 King, C. & Sprent, J. Dual Nature of Type I Interferons in SARS-CoV-2-Induced Inflammation. Trends Immunol 42, 312–322, doi:10.1016/j.it.2021.02.003 (2021).

8 Ziegler, C. G. K. et al. SARS-CoV-2 Receptor ACE2 Is an Interferon-Stimulated Gene in Human Airway Epithelial Cells and Is Detected in Specific Cell Subsets across Tissues. Cell 181, 1016–1035 e1019, doi:10.1016/j.cell.2020.04.035 (2020).

9 Britten, R. J. & Davidson, E. H. Gene regulation for higher cells: a theory. Science (New York, N.Y.) 165, 349–357 (1969).

10 Hale, B. G. Antiviral immunity triggered by infection-induced host transposable elements. Curr Opin Virol 52, 211–216, doi:10.1016/j.coviro.2021.12.006 (2021).

11 Chuong, E. B., Elde, N. C. & Feschotte, C. Regulatory evolution of innate immunity through co-option of endogenous retroviruses. Science (New York, N.Y.) 351, 1083–1087, doi:10.1126/science.aad5497 (2016).

12 Wang, M., Wang, L. Y., Liu, H. Z., Chen, J. J. & Liu, D. Transcriptome Analyses Implicate Endogenous Retroviruses Involved in the Host Antiviral Immune System through the Interferon Pathway. Virol Sin 36, 1315–1326, doi:10.1007/s12250-021-00370-2 (2021).

13 Ye, M. et al. Specific subfamilies of transposable elements contribute to different domains of T lymphocyte enhancers. Proc Natl Acad Sci USA 117, 7905–7916, doi:10.1073/pnas.1912008117 (2020).

14 Macchietto, M. G., Langlois, R. A. & Shen, S. S. Virus-induced transposable element expression up-regulation in human and mouse host cells. Life science alliance 3, doi:10.26508/lsa.201900536 (2020).

15 Tunbak, H. et al. The HUSH complex is a gatekeeper of type I interferon through epigenetic regulation of LINE-1s. Nat Commun 11, 5387, doi:10.1038/s41467-020-19170-5 (2020).

16 Krischuns, T. et al. Phosphorylation of TRIM28 Enhances the Expression of IFN-β and Proinflammatory Cytokines During HPAIV Infection of Human Lung Epithelial Cells. Front Immunol 9, doi:10.3389/fimmu.2018.02229 (2018).

17 Chiappinelli, K. B. et al. Inhibiting DNA Methylation Causes an Interferon Response in Cancer via dsRNA Including Endogenous Retroviruses. Cell 162, 974–986, doi:10.1016/j.cell.2015.07.011 (2015).

18 Gazquez-Gutierrez, A., Witteveldt, J., Heras, S. R. & Macias, S. Sensing of transposable elements by the antiviral innate immune system. Rna 27, 735–752, doi:10.1261/rna.078721.121 (2021).

19 Pehrsson, E. C., Choudhary, M. N., Sundaram, V. & Wang, T. The epigenomic landscape of transposable elements across normal human development and anatomy. Nature Communications 10, 1–16 (2019).

20 Bogu, G. K., Reverter, F., Marti-Renom, M. A.,, Snyder, M. P. & Guigó, R. Atlas of transcriptionally active transposable elements in human adult tissues. BioRxiv, 714212 (2019).

21 Franceschi, C., Garagnani, P., Parini, P., Giuliani, C. & Santoro, A. Inflammaging: a new immune-metabolic viewpoint for age-related diseases. Nat Rev Endocrinol 14, 576–590, doi:10.1038/s41574-018-0059-4 (2018).

22 Zhang, W. et al. A Werner syndrome stem cell model unveils heterochromatin alterations as a driver of human aging. Science 348, 1160–1163 (2015).

23 Perez, R. F., Tejedor, J. R., Bayon, G. F., Fernández, A. F. & Fraga, M. F. Distinct chromatin signatures of DNA hypomethylation in aging and cancer. Aging Cell 17, e12744 (2018).

24 Galani, I.-E. et al. Untuned antiviral immunity in COVID-19 revealed by temporal type I/III interferon patterns and flu comparison. Nat Immunol, 1–9 (2020).

25 Tokuyama, M. et al. ERVmap analysis reveals genome-wide transcription of human endogenous retroviruses. Proc Natl Acad Sci USA 115, 12565–12572, doi:10.1073/pnas.1814589115 (2018).

26 Karaderi, T. et al. Host Genetics at the Intersection of Autoimmunity and COVID-19: A Potential Key for Heterogeneous COVID-19 Severity. Front Immunol 11, doi:ARTN58611110.3389/fimmu.2020.586111 (2020).

27 Ferrarini, M. G. et al. Genome-wide bioinformatic analyses predict key host and viral factors in SARS-CoV-2 pathogenesis. Commun Biol 4, 590, doi:10.1038/s42003-021-02095-0 (2021).

28 Marston, J. L. et al. SARS-CoV-2 infection mediates differential expression of human endogenous retroviruses and long interspersed nuclear elements. JCI Insight 6, doi:10.1172/jci.insight.147170 (2021).

29 Rebendenne, A. et al. SARS-CoV-2 triggers an MDA-5-dependent interferon response which is unable to control replication in lung epithelial cells. J Virol, doi:10.1128/JVI.02415-20 (2021).

30 Schmidt, N. et al. An influenza virus-triggered SUMO switch orchestrates co-opted endogenous retroviruses to stimulate host antiviral immunity. Proc Natl Acad Sci USA 116, 17399–17408, doi:10.1073/pnas.1907031116 (2019).

31 Takayama, K. In Vitro and Animal Models for SARS-CoV-2 research. Trends Pharmacol Sci 41, 513–517, doi:10.1016/j.tips.2020.05.005 (2020).

32 Badarinarayan, S. S. & Sauter, D. Switching Sides: How Endogenous Retroviruses Protect Us from Viral Infections. Journal of Virology 95, doi:ARTNe02299-2010.1128/JVI.02299-20 (2021).

33 Roulois, D. et al. DNA-Demethylating Agents Target Colorectal Cancer Cells by Inducing Viral Mimicry by Endogenous Transcripts. Cell 162, 961–973, doi:10.1016/j.cell.2015.07.056 (2015).

34 Davis, M. I. et al. Comprehensive analysis of kinase inhibitor selectivity. Nat. Biotechnol. 29, 1046–U1124, doi:10.1038/nbt.1990 (2011).

35 Enriquez-Gasca, R., Gould, P. A. & Rowe, H. M. Host Gene Regulation by Transposable Elements: The New, the Old and the Ugly. Viruses 12, doi:10.3390/v12101089 (2020).

36 Chuong, E. B., Elde, N. C. & Feschotte, C. Regulatory activities of transposable elements: from conflicts to benefits. Nat Rev Genet 18, 71–86, doi:10.1038/nrg.2016.139 (2017).

37 Schreiber, J., Bilmes, J. & Noble, W. S. Completing the ENCODE3 compendium yields accurate imputations across a variety of assays and human biosamples. Genome Biol 21, 82, doi:10.1186/s13059-020-01978-5 (2020).

38 Sorek, M., Cohen, L. R. Z. & Meshorer, E. Open chromatin structure in PolyQ disease-related genes: a potential mechanism for CAG repeat expansion in the normal human population. NAR Genom Bioinform 1, e3, doi:10.1093/nargab/lqz003 (2019).

39 Chung, N. et al. Transcriptome analyses of tumor-adjacent somatic tissues reveal genes co-expressed with transposable elements. Mob DNA 10, 39, doi:10.1186/s13100-019-0180-5 (2019).

40 Medvedeva, Y. A. et al. EpiFactors: a comprehensive database of human epigenetic factors and complexes. Database (Oxford) 2015, bav067, doi:10.1093/database/bav067 (2015).

41 Chen, K. et al. Methyltransferase SETD2-Mediated Methylation of STAT1 Is Critical for Interferon Antiviral Activity. Cell 170, 492–506 e414, doi:10.1016/j.cell.2017.06.042 (2017).

42 Li, F. et al. The histone mark H3K36me3 regulates human DNA mismatch repair through its interaction with MutSalpha. Cell 153, 590–600, doi:10.1016/j.cell.2013.03.025 (2013).

43 Dhayalan, A. et al. The Dnmt3a PWWP domain reads histone 3 lysine 36 trimethylation and guides DNA methylation. J Biol Chem 285, 26114–26120, doi:10.1074/jbc.M109.089433 (2010).

44 Li, X. et al. The MOV10 helicase inhibits LINE-1 mobility. J Biol Chem 288, 21148–21160, doi:10.1074/jbc.M113.465856 (2013).

45 De Cecco, M. et al. L1 drives IFN in senescent cells and promotes age-associated inflammation. Nature 566, 73–78, doi:10.1038/s41586-018-0784-9 (2019).

46 Barral, A. et al. SETDB1/NSD-dependent H3K9me3/H3K36me3 dual heterochromatin maintains gene expression profiles by bookmarking poised enhancers. Mol Cell 82, 816–832 e812, doi:10.1016/j.molcel.2021.12.037 (2022).

47 Burns, K. H. Our Conflict with Transposable Elements and Its Implications for Human Disease. Annu Rev Pathol 15, 51–70, doi:10.1146/annurev-pathmechdis-012419-032633 (2020).

48 Bolger, A. M., Lohse, M. & Usadel, B. Trimmomatic: a flexible trimmer for Illumina sequence data. Bioinformatics 30, 2114–2120, doi:10.1093/bioinformatics/btu170 (2014).

49 Dobin, A. et al. STAR: ultrafast universal RNA-seq aligner. Bioinformatics 29, 15–21, doi:10.1093/bioinformatics/bts635 (2013).

50 Jin, Y., Tam, O. H., Paniagua, E. & Hammell, M. TEtranscripts: a package for including transposable elements in differential expression analysis of RNA-seq datasets. Bioinformatics 31, 3593–3599, doi:10.1093/bioinformatics/btv422 (2015).

51 Love, M. I., Huber, W. & Anders, S. Moderated estimation of fold change and dispersion for RNA-seq data with DESeq2. Genome Biol. 15, 550, doi:10.1186/s13059-014-0550-8 (2014).

52 Liao, Y., Smyth, G. K. & Shi, W. featureCounts: an efficient general purpose program for assigning sequence reads to genomic features. Bioinformatics 30, 923–930, doi:10.1093/bioinformatics/btt656 (2014).

53 Yin, T., Cook, D. & Lawrence, M. ggbio: an R package for extending the grammar of graphics for genomic data. Genome Biol 13, R77, doi:10.1186/gb-2012-13-8-r77 (2012).

54 Stukalov, A. et al. Multilevel proteomics reveals host perturbations by SARS-CoV-2 and SARS-CoV. Nature 594, 246–252, doi:10.1038/s41586-021-03493-4 (2021).

55 Samavarchi-Tehrani, P. et al. A SARS-CoV-2 – host proximity interactome. bioRxiv, 2020.2009.2003.282103, doi:10.1101/2020.09.03.282103 (2020).

56 Laurent, E. M. N. et al. Global BioID-based SARS-CoV-2 proteins proximal interactome unveils novel ties between viral polypeptides and host factors involved in multiple COVID19-associated mechanisms. bioRxiv, 2020.2008.2028.272955, doi:10.1101/2020.08.28.272955 (2020).

57 Gordon, D. E. et al. Comparative host-coronavirus protein interaction networks reveal pan-viral disease mechanisms. Science 370, doi:10.1126/science.abe9403 (2020).

58 Gordon, D. E. et al. A SARS-CoV-2 protein interaction map reveals targets for drug repurposing. Nature 583, 459–468, doi:10.1038/s41586-020-2286-9 (2020).

59 Wang, L. et al. Comparative influenza protein interactomes identify the role of plakophilin 2 in virus restriction. Nat Commun 8, 13876, doi:10.1038/ncomms13876 (2017).

60 Calderwood, M. A. et al. Epstein-Barr virus and virus human protein interaction maps. Proc Natl Acad Sci U S A 104, 7606–7611, doi:10.1073/pnas.0702332104 (2007).

61 Li, M. et al. Identification of antiviral roles for the exon-junction complex and nonsense-mediated decay in flaviviral infection. Nat Microbiol 4, 985–995, doi:10.1038/s41564-019-0375-z (2019).

