## Supplementary figures and images for "Impaired activation of Transposable Elements in SARS-CoV-2 infection"

### Supplemental Figure 1

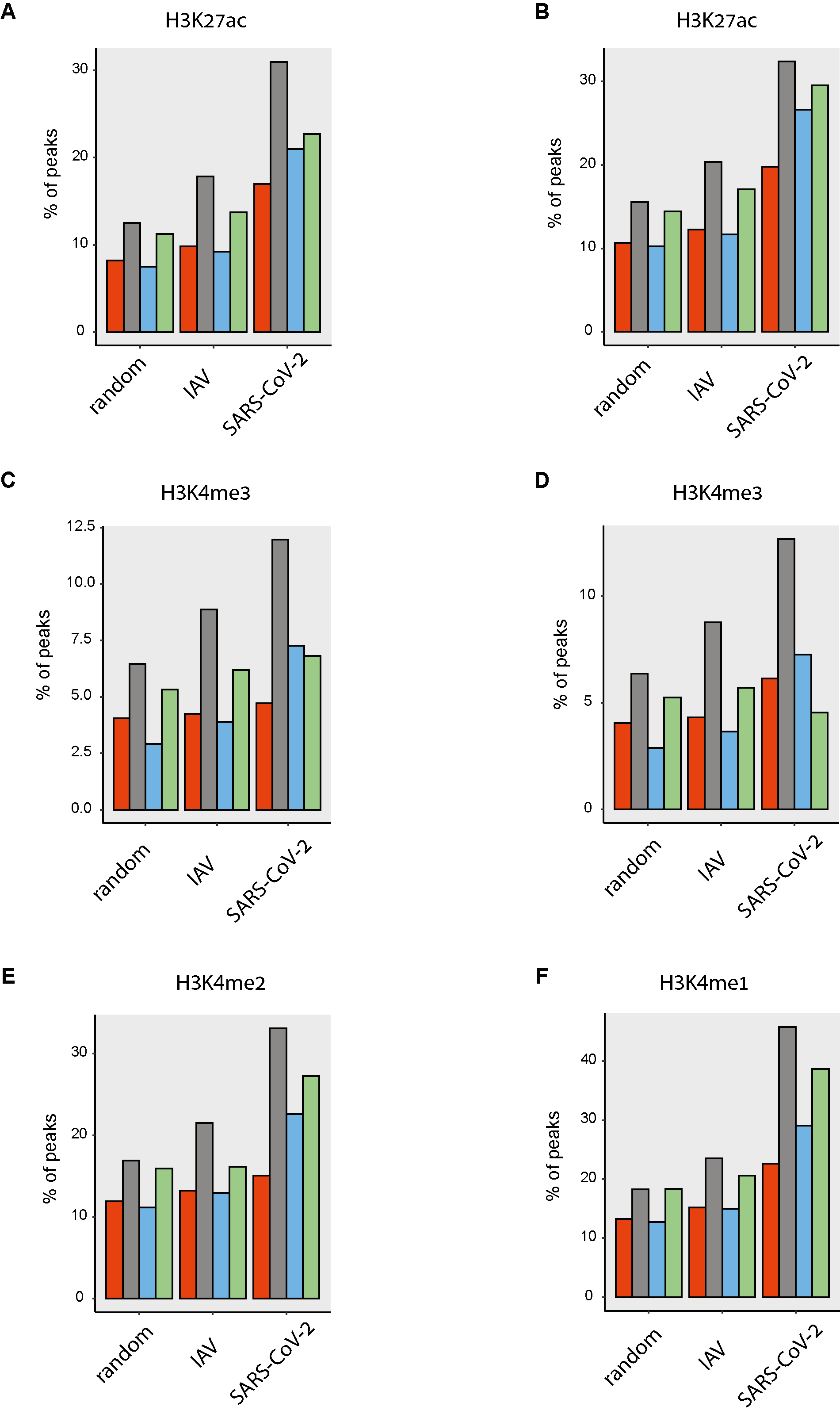

### Supplemental Figure 2

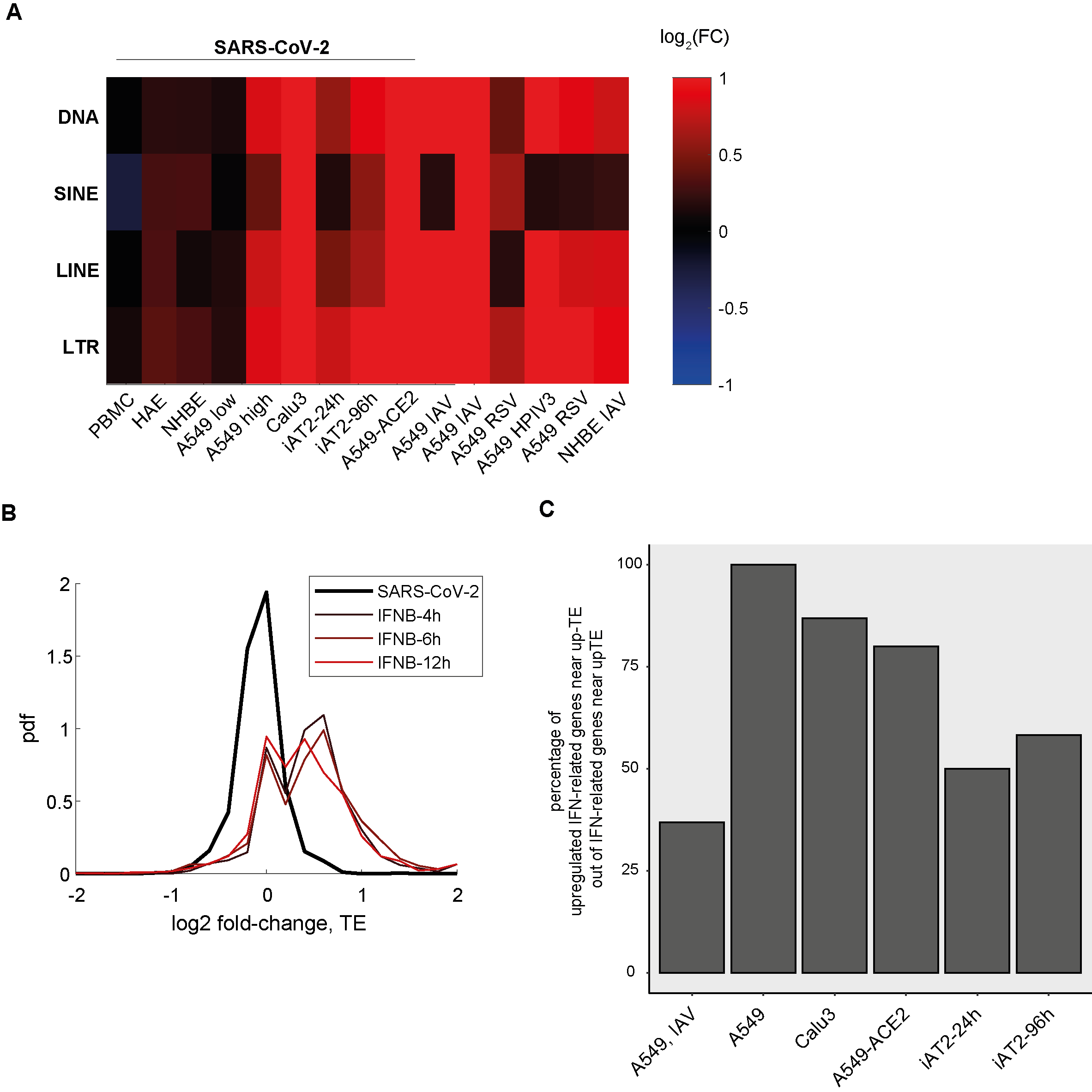

### Supplemental Figure 3

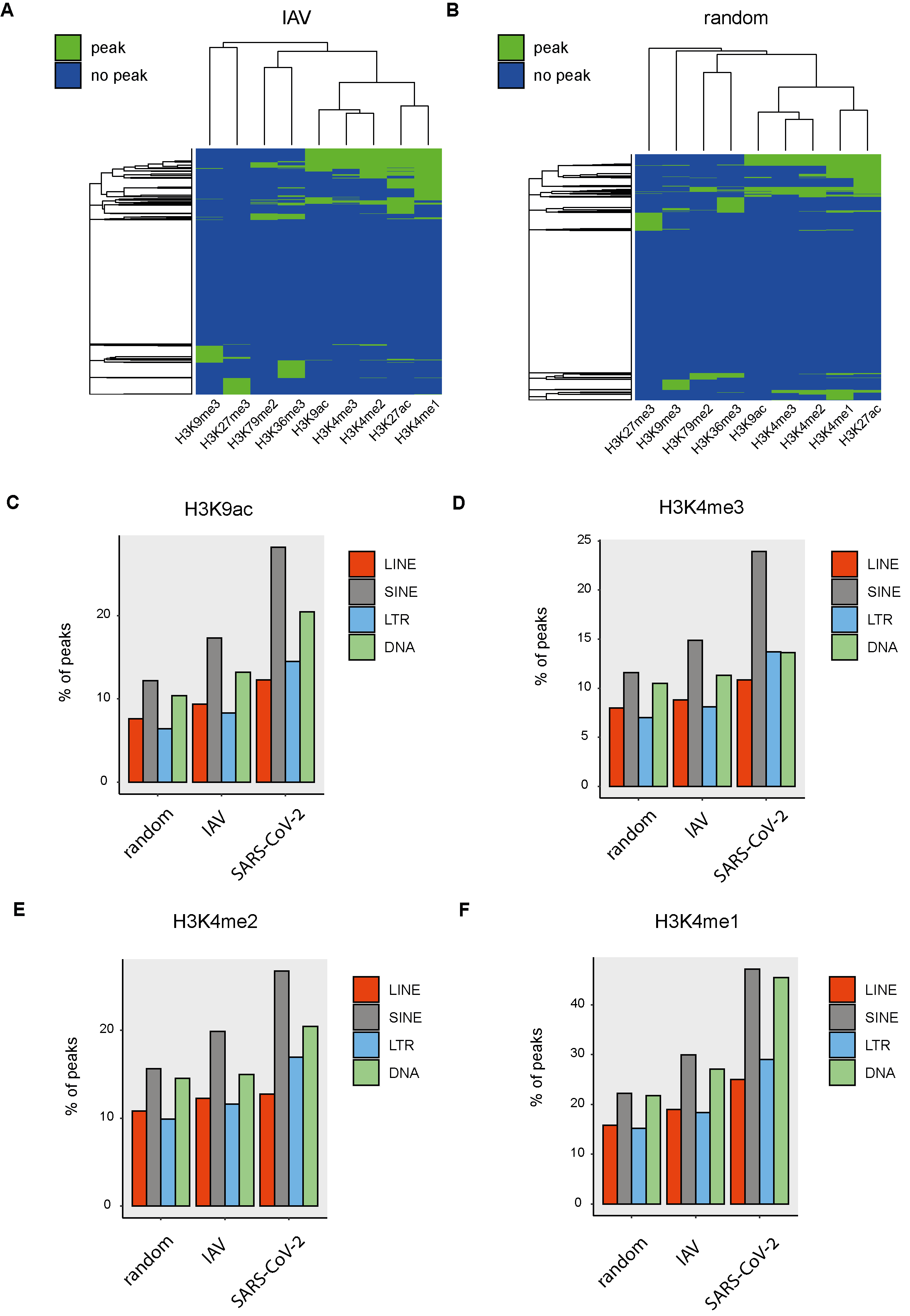
